# cinaR: A comprehensive R package for the differential analyses and functional interpretation of ATAC-seq data

**DOI:** 10.1101/2021.03.05.434143

**Authors:** E Onur Karakaslar, Duygu Ucar

## Abstract

**Summary:** ATAC-seq is a frequently used assay to study chromatin accessibility levels. Differential chromatin accessibility analyses between biological groups and functional interpretation of these differential regions are essential in ATAC-seq data analyses. Although distinct methods and analyses pipelines are developed for this purpose, a stand-alone R package that combines state-of-the art differential and functional enrichment analyses pipelines is missing. To fill this gap, we developed *cinaR (***C**hromat<u>**in A**nalyses in **R**</u>*)*, which is a single wrapper function and provides users with various data analyses and visualization options, including functional enrichment analyses with gene sets curated from multiple sources.

**Availability and implementation:** *cinaR* is an R/CRAN package which is under GPL-3 License and its source code is freely accessible at https://CRAN.R-project.org/package=cinaR.

Gene sets are available at https://CRAN.R-project.org/package=cinaRgenesets.

Bone marrow ATAC-seq data is available at https://www.ncbi.nlm.nih.gov/geo/query/acc.cgi?acc=GSE165120

## 1 Introduction

Assay for transposase accessible chromatin with high-throughput sequencing (ATAC-seq) is a technology for probing the chromatin-accessibility levels from small cell numbers (Buenrostro et al., 2013). Briefly, Tn5 transposase cuts the open chromatin regions (OCRs); these fragments are sequenced using high-throughput sequencing and then aligned to the genome to uncover ATAC-seq peaks mapping to OCRs (Tsompana and Buck., 2014). ATAC-seq is highly adopted by the scientific community including its application to study single cell epigenomes (Chung et al. 2019, Zhang et al., 2021, Satpathy et al. 2019).

ATAC-seq data analyses guidelines have been developed including by the ENCODE project (ATAC-seq Data Standards and Processing Pipeline – ENCODE, 2020) and others (Gaspar, 2020). However, an easy-to-use R package for this purpose is missing. To fill this gap, we developed, *cinaR*, (***C**hromat<u>**in A**nalyses in **R**</u>*), which can conduct differential accessibility analyses, batch correction, and functional enrichment of differential peak results. *cinaR* accomplishes these within a single wrapper function in order to provide an easy-to-use interface for users while maintaining high customizability *via* various options for data analyses and visualization. To complement functional enrichments in *cinaR*, we also implemented an additional CRAN/R package which contains gene sets that are carefully curated from different sources especially for the analyses of immune cells (https://github.com/eonurk/cinaR-genesets).

## 2 Materials and Methods

Starting from a consensus peaks matrix, *cinaR* filters the peaks, annotates them to their corresponding genes, conducts differential and functional enrichment analyses with customizable options and then let users to visualize their findings (summarized in **Figure 1A**).

**Figure 1.**
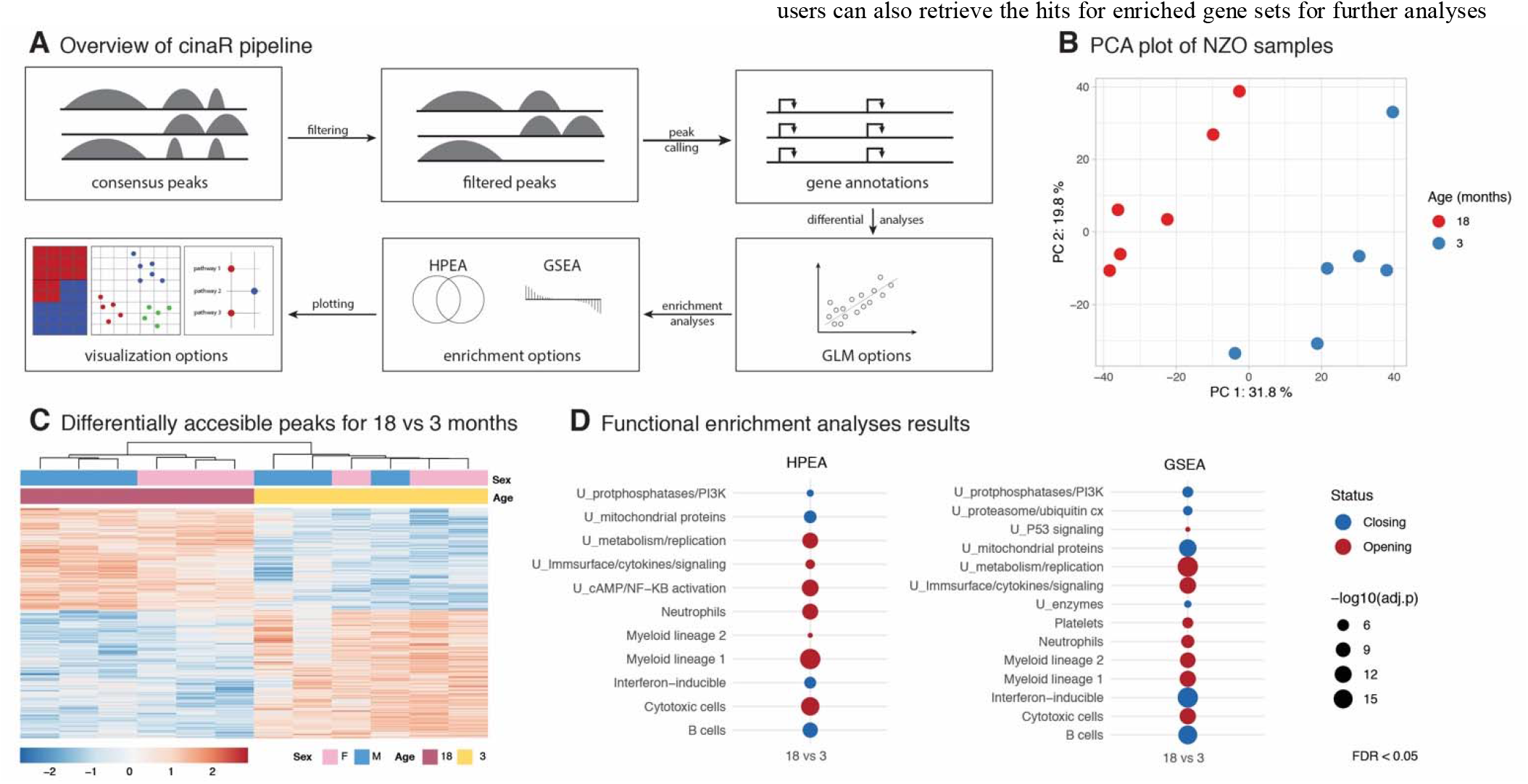
(**A**) Overall workflow schematic of the *cinaR* pipeline. (**B**) PCA plot clearly separates 3 months (blue) and 18 months (red) NZO mice samples using filtered and normalized peaks (n=34116). (**C**) Heatmap of Differentially Accessible (DA) peaks at FDR=0.05. In total there are 6653 peaks (2956 opening, 3697 closing with age). (**D**) Functional enrichment analyses of DA peaks using HPEA and GSEA options. It yielded similar results most notably regarding the up-regulation of pro-inflammatory pathways such as Myeloid lineage 1,2 and NfKB activation.

### 2.1 Implementation Details

#### 2.1.1 Peak filtering and annotation to genes

*cinaR* requires a consensus peak matrix and a vector that indicates the biological/clinical grouping of the ATAC-seq samples to be used in the differential analyses. First, ATAC-seq peaks are kept for down-stream analyses if the count-per-million (CPM) normalized counts are above a certain threshold (>0.5) for more than k samples (default k=2). Then these selected peaks are annotated to the closest gene based on distance to Transcription Start Site (TSS) using *ChIPseeker* (Yu et al, 2015). The annotated peaks are further filtered with a threshold (default 50Kb) of their absolute distance to transcription start sites (TSS).

#### 2.1.2 Differ ential accessibility analyses

To conduct differential accessibility analyses, a design matrix is built to conduct pairwise comparisons among all distinct biological/clinical groups provided by the user. For n groups, 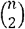 comparisons are conducted. Users can select among four alternative methods for differential analyses: *edgeR* (Robinson et al, 2009), *limma-voom* and *limma-trend* (Richie et al, 2015) and *DESeq2* (Love et al, 2014). *edgeR* is selected as the default option with FDR = 0.05. If the input consensus peak matrix is composed of raw counts (e.g., CPM), we suggest using either *edgeR* or *DESeq2*. If the library sizes are heterogenous *limma-voom* is recommended. If the consensus peaks are already normalized, *limmatrend* should be used. To eliminate potential batch-effects, we implemented two alternative methods. If batch information is not provided by the user, surrogate variable analyses (SVA) is conducted to detect unknown batch effects (Leek and Storey, 2020). This option will calculate the number surrogate variables (SVs) automatically and add these to the design matrix. Users also have an option for using a certain number of SVs instead of all significant ones. On the other hand, if the batches are known, the batch information is included in the design matrix of the linear model and used as a covariate in the differential analyses.

#### 2.1.3 Functional enrichment analyses

For functional enrichment analyses, *cinaR* provides two options: hypergeometric p-value (HPEA) and gene set enrichment analyses (GSEA) (Subramanian et al., 2005). If HPEA is selected, the differential peaks are split based on the direction of changes (opening versus closing peaks) and enrichment p-values are calculated for each group separately. If GSEA is selected, annotated peaks are sorted with respect to their fold change. If a gene is associated with multiple peaks, the one that is closest to the gene TSS is used in these analyses. For both methods, the enrichment p-values are corrected using Benjamini-Hochberg procedure (Hochberg and Benjamini, 1990) and adjusted p-values are reported. The and interpretation of the data. *cinaR* supports two human (hg19 and hg38) and one mouse genome (mm10) versions. The default option is hg38, yet if not set by the user, it will throw a warning to avoid genome mismatching.

#### 2.2 Gene sets curated for *cinaR*

Functional enrichment of differential peaks is an important yet daunting task. We have curated several gene sets from multiple sources throughout years for this purpose, which are provided within another CRAN/R package that we use as part of the *cinaR* pipeline (https://CRAN.R-project.org/package=cinaR). This includes six different gene sets that are particularly effective for the study of immune cells: immune modules, PBMC-scRNAseq, wikipathways, wikipathway-inflammation, activated-immune gene sets, and gene sets from the DICE project (dice-major). Immune modules are a total of 28 gene sets that are compiled from gene expression profiles of human peripheral blood mononuclear cells (PBMCs) samples including from healthy and diseased samples (Chaussabel *et al*., 2008). PBMC-scRNAseq consists of 15 modules, where each gene set represents cell type specific genes for immune cell subsets within PBMCs, that are inferred from single cell RNA-seq PBMC data (Márquez et al., 2020; Nehar-Belaid et al., 2020). WikiPathways are 671 biologically meaningful pathways which are created by a community-based collaborative effort. In addition to all WikiPathways, we also curated a subset of these (n=50) that are inflammation and immune system related and labeled them as Wikipathways-inflammation (Pico *et al*., 2008). Lastly, we curated 6 modules from the dice database which includes the transcriptional signatures of different immune cell types (Schmiedel *et al*., 2018). This additional package is also freely accessible under GPL-3 license at https://cran.r-project.org/package=cinaRgenesets. In addition, users can also incorporate their own gene sets into the *cinaR* pipeline by using .*gmt* format, which provides extra flexibility for functional enrichment analyses.

## 3 Results

To benchmark *cinaR*, we generated ATAC-seq data from bone-marrow (GEO accession GSE165120) in short-lived NZO/HILtJ (NZO) mice strain at two age groups: 3-month-old (n=6, young) and 18-month-old (n=6, old) animals. These samples were analyzed using *cinaR* to identify age related changes in the chromatin accessibility maps. **Figure 1B** shows the PCA plot for filtered and normalized peaks (n=34116), where samples are separated with respect to age (no batch effects are detected). Using default settings of *cinaR*, we identified 6653 differentially accessible peaks between young and old animals (2956 opening, 3697 closing with age) at FDR 5%. Functional enrichment analyses of these peaks using HPEA and GSEA options with the immune modules revealed that as expected pro-inflammatory modules are activated (myeloid lineage, NFkB) with age (**Figure 1C**).

## 4 Discussion

ATAC-seq is a widely used technology to study open chomatin regions in the genome. Although there are distinct pipelines for differential and functional enrichment analyses of ATAC-seq data, a pipeline which combines the state-of-art methodologies is still missing. Here, we presented *cinaR*, a CRAN/R package that provides users with flexibility to run both analyses with their methods of choice. In addition to that it has options to correct for batch effects as well as covariates, and it also includes highly customizable functions for visualizations. We also implemented another CRAN/R package (*cinaR-genesets*) along with the original one where we shared immune-related gene sets that we have curated from different resources.

